# Genomic sampling and population structure of farmer-maintained varieties reveal previously uncharacterized diversity of *Theobroma cacao* L. in Costa Rica

**DOI:** 10.64898/2026.03.30.715340

**Authors:** Emma M. Herrighty, Chelsea D Specht, Michael A. Gore, Luis Solano, Joyce Estrada-Gamboa, Carlos Eduardo Hernandez, Hale Ann Tufan, Jacob B. Landis

## Abstract

Understanding crop genetic diversity is essential for conservation and breeding, yet farmer-maintained germplasm remains largely underrepresented in genomic studies. *Theobroma cacao* L. has a complex domestication history and extensive global diversity, and cacao currently cultivated in Central America, particularly in Costa Rica, has been understudied compared to South American and Mexican cultivars despite cultural and historical importance. In this study, we investigate the genetic diversity of cacao from farmer-managed systems across Costa Rica to search for Criollo germplasm and identify and characterize any unique local genetic groups. Ninety-four trees were sampled from 17 farms across four regions of the country and sequenced using whole genome resequencing. Farmer materials were analyzed alongside 166 previously characterized reference accessions representing major cacao genetic groups. Population structure analyses, phylogenetic reconstruction, and network approaches revealed that Costa Rican cacao encompasses multiple known genetic groups, including Criollo-derived lineages, while also harboring locally distinct diversity not fully represented in current global reference collections. Analyses revealed close kinship between many accessions with no clear geographic patterns corresponding to the observed population differentiation, reflecting the effects of farmers in creating dominant patterns of gene flow through seed-saving, clonal propagation, and sharing genotypes among farms. Heterozygosity levels varied substantially among individuals, consistent with a mixture of highly inbred Criollo trees and more heterozygous, admixed genotypes. We find that farmer-managed cacao systems are reservoirs of genetic diversity, including possibly rare or historically important lineages, underscoring the value of these farming systems for effective conservation and management of genomic resources for cacao resilience and improvement.

## 1. Introduction

Global crop diversity underpins food security, climate resilience, and sustainable agricultural systems, yet it remains unevenly characterized and therefore underutilized across many crop species (Khoury et al., 2022). While genomic resources often incorporate cultivated and wild accessions, the deliberate prioritization of landraces and traditional planting materials has been less common and is highly crop dependent (Ramirez-Villegas et al., 2022; Lazaridi et al., 2024). Research on neglected crops such as sesame (Yu et al., 2019) and watermelon (Sun et al., 2025) have included multiple landrace accessions, while the tomato pangenome includes much of the vast diversity of heirloom cultivars (Gao et al., 2019). A genetic characterization of farmer-managed germplasm can highlight the hidden diversity maintained within crop systems, especially when cultivars have been developed over many generations and across heterogeneous landscapes (Brush, 2000; Brusse et al., 2024). Farming systems and their locally-adapted varieties represent dynamic reservoirs of genetic variation shaped by environmental pressures and cultural practices (Enjalbert et al., 2011; Shrestha, Gudino & Angeles-Shim, 2025).

*Theobroma cacao* L. (Malvaceae) is a diploid tree species (2n = 2x = 20) and the source of cocoa, a historically, culturally and economically important tropical crop (reviewed in Young, 1994; Bartley, 2005). While the Amazon basin is considered the center of origin for cacao, a complex domestication history, along with both natural and human-mediated dispersals, have spread cacao throughout Central and South America (Motamayor et al., 2002; Piperno, 2011; Pickersgill, 2016; Cornejo et al., 2018; Lanaud et al., 2024). Archaeological evidence indicates that cacao has been cultivated for over 5,000 years (Lanaud et al., 2024). Traditional classifications distinguish Criollo, Forastero, and Trinitario types; however, advances in genetic studies have revealed more than 10 major genetic groups (Motamayor et al., 2008; Zhang et al., 2012; Lachenaud & Motamayor, 2017, Fouet et al., 2022; Argout et al., 2023).

The global genetic diversity of cacao has revealed numerous genetic clusters shaped by periods of geographic isolation and subsequent diversification (Motamayor et al., 2008; Cornejo et al., 2018; Argout et al., 2023), yet the true diversity of the species remains unclear (Lanaud, Motamayor, & Sounigo, 2003; Motamayor et al., 2008; Bekele and Phillips-Mora, 2019). This knowledge gap has implications for breeding, as well as for effective long-term conservation of cacao genetic resources. Effective germplasm conservation depends on materials being well-characterized, documented, and representative of the species’ full geographic and cultural range (Schreiber et al., 2024; Stange, Barrett, & Hendry, 2021). While significant collection efforts have been made, especially the center of origin, the relationships between conserved accessions and cultivated cacao remains poorly resolved, limiting both conservation value and contributions to the genetic improvement of cacao (Cornejo et al., 2018; González-Orozco, Osorio-Guarín &Yockteng, 2022; Todd et al., 2025).

Previous genetic studies in cacao (Zhang et al., 2012; Fouet et al. 2022) and the recent pangenome (Argout et al., 2023) underrepresent cacao-producing regions such as Central America and specifically Costa Rica. Cacao has longstanding historical, cultural, and economic importance to Costa Rica, remaining deeply tied to Indigenous communities and heritage production systems (Oviedo y Valdes, 1853; Kaviany, 2020; Steinbrenner et al., 2021). Existing heritage and locally-managed cacao accessions are likely sources of unique genetic variation absent from current reference collections (Lavoie, Thomas & Olivier, 2023; Todd et al., 2025). Knowledge of local diversity, particularly heritage trees, is crucial for sustaining regional production, contributing to *in situ* conservation, and identifying alleles for breeding programs (Ji et al., 2013; Vázquez-Ovando et al., 2014). Characterizing this diversity should be a priority since local materials are increasingly under threat of being displaced by globally-sourced “improved” cultivars (Bartley, 2005; Bidot Martinez et al., 2015).

Criollo cacao exemplifies both the value and vulnerability of localized diversity. Historically prized for its superior flavor, production of Criollo had apparently declined so dramatically that Braudeau (1969) described the variety as virtually no longer cultivated. Disease susceptibility, genetic erosion by hybridization, and the mass replacement of older plantings with improved clones raised concerns about the presence and diversity of the ancient Criollo lineage. A recent distinction between “Ancient” (genetically pure, or unadmixed) Criollo and “Modern” (admixed) Criollo (reviewed by Lachenaud and Motanayor, 2017) ultimately underscores the recognition of local gene flow and farmer selection in reshaping the genetic makeup of this lineage. Modern Criollos are more common today and remain closely related to Ancient Criollo; their persistence as local cultivars demonstrates the potential for new genetic subgroups to arise from natural or managed gene flow between Ancient Criollo and local or introduced cacao lineages (Bidot Martinez et al., 2015). Despite narratives of the rarity of Criollo germplasm, Ancient Criollo trees are believed to still exist in farmer plots and in the wild, conserved and disseminated by producers who have attachments to the regional historic, cultural, and production value of these materials (Lachenaud & Motamayor, 2017). Ancient Criollo trees continue to be sought out as relics for their sociocultural value and ties to local tradition; while also serving as reservoirs of genes and alleles with potential value for breeding and selection programs focused on local adaptation and disease resistance (Motilal et al., 2010; Vázquez-Ovando et al., 2014).

To elucidate the genetic diversity of Costa Rican cacao and investigate the presence and prevalence of Criollo germplasm, we sampled older, heritage, and non-improved plantings from select local and small farms across the country. Farmers were asked to identify trees that they believed had local and/or historic ties to the area. These materials were sampled and genetically characterized via whole genome resequencing, enabling high-resolution analyses of population structure, admixture patterns, levels of heterozygosity, and distinctiveness of a diversity of Criollo-derived lineages. The specific objectives of this study are to (a) identify the major genetic groups represented in farmers’ fields based on known references, in particular trees reported to have Criollo origins, (b) identify and characterize genetic groups unique to Costa Rica, and (c) understand patterns of genetic admixture across sampled populations.

## 2. Materials and Methods

### 2.1 Plant material

Sampling sites were selected based on recommendations from leaders within local cacao grower organizations, "las plataformas cacao." Growers were included in the study if they self-identified or were believed to steward local cacao germplasm instead of growing primarily genetically-improved, clonally-propagated cultivars. Between October and December 2024, 94 cacao trees were sampled from 17 farms spanning four regions of Costa Rica (Figure 1). From north to south, these regions were Guatuso-Upala (GTS-UPA), Guácimo (GCM), Talamanca (TAL), and Puntarenas (PUNT). Farms were assigned an abbreviated region code and each tree sampled from that farm was numbered such that the sample code reflects Region (CODE), Farm (#), and Tree ID (-#) (*e.g*., UPA1-1).

**Figure 1.**
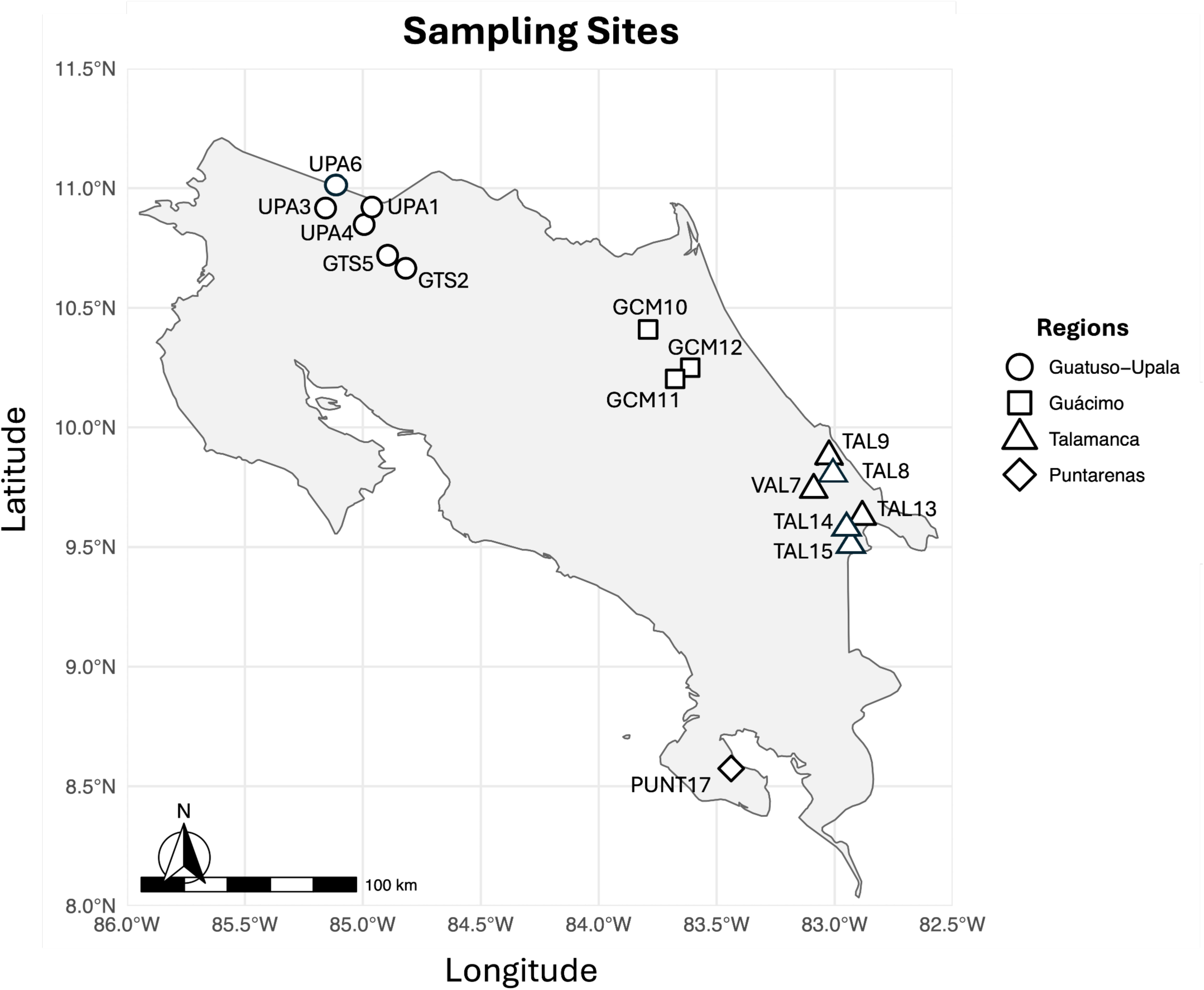
Map of the 17 farms where samples were collected across four designated Regions (see key). Each point represents one farm, with distinct shapes indicating the designated Region. Farms were assigned an abbreviated region code (Guatuso = GTS; Upala = UPA; Guácimo = GCM; Talamanca = TAL; Puntarenas = PUNT), followed by a number indicating the order in which collections were made.

Within each farm, trees were selected based on farmers identifying trees they believed to be locally adapted or Criollo. Provenance or genetic background of the material was recorded when known. Farmers were asked to identify unique and valued traits (*i.e.*, pod shape and color, productivity, flavor profile) as a means of including morphological and genetic diversity.

Latitude and longitude of each tree were recorded at the time of collection. Three young, mature leaves were collected per tree and stored in silica gel prior to high-throughput magnetic bead-based DNA extraction performed by Intertek (Sweden). DNA concentration was quantified using an Invitrogen Qubit™ Fluorometer (Carlsbad, California, USA) with the dsDNA High Sensitivity assay using 2 µl.

### 2.2 Library Preparation and Sequencing

Genome resequencing libraries were constructed using a Nextera tagmentation protocol optimized by the Cornell Biotechnology Resource Center (BRC) Genomics Innovation Hub. This protocol is similar to published protocols such as Hennig et al., (2018) and Vonesch et al., (2021), as well as those discussed in Stewart et al., (2024), but uses a homebrewed Tn5.3 transposome. The tagmentation-based protocol used 100 ng of normalized genomic DNA, 3 µL of 2 mM transposome, a 10-minute tagmentation incubation step at 55℃, a 7-minute incubation with 3 µL of 2% SDS at 55℃ to stop the tagmentation reaction, with a final addition of 3 µL of 10% triton. A bead cleanup using homemade cleaning beads (Rowan et al., 2017) was done with 0.8x volume of beads. Libraries were amplified and barcodes added using 10 cycles of PCR with Nextera UDI primers (initial five minute extension at 72℃, with denaturation at 98℃ for 30 seconds, followed by 10 cycles of 98℃ for 10 sec, 67℃ for 30 seconds, and 72℃ for one minute, with a final extension at 72℃ for five minutes). Eight libraries were verified by fragment size analysis using an Agliant 5200 Fragment Analyzer to ensure that most of the DNA fragments were between 450 and 800 bp prior to pooling. All samples were pooled by equal mass targeting 30 ng per sample, and the resulting pool was bead cleaned with sparQ beads (Quanta bio, Beverley, MA, USA) to concentrate the volume prior to size selection of 450 and 800 bp with the pippin at the Cornell BRC Genomics Facility. Size selected pools were sequenced on one lane of an Illumina NovaSeqX 25B (2 x 150).

### 2.3 Reference Accessions

A reference panel of 166 previously published *Theobroma* accessions spanning the geographic and genetic diversity of cacao (Supplemental data S2) were selected for comparative analyses representing the 10 genetic groups identified by Motamayor et al., (2008) plus a total of five additional groups described by Fouet et al., (2022) and Argout et al., (2023). Each accession was retrieved from the NCBI Sequence Read Archive using the SRA Toolkit v3.0.10 (SRA Toolkit Development Team) and processed concurrently with the newly sequenced samples.

### 2.4 Bioinformatic processing

Two data sets were constructed to explore the existing genetic diversity of Costa Rican cacao. (A) The first included samples from all 15 reference groups plus our Costa Rican collections (n = 259) and was used primarily for population structure analyses to assess how newly sampled materials fit with previously defined cacao genetic groups. (B) The second consisted of only our Costa Rican collections (n = 93) and was used for further analyses investigating patterns of gene flow and genetic diversity across the sampled farms.

Sequencing reads were cleaned with fastp v0.23.4 (Chen, 2023) to filter out low quality bases (Phred scores < 20), remove PolyG strings, and discard reads shorter than 75 bp. Cleaned reads were mapped to the B97-61/B2 Criollo cacao reference genome (v2) (Argout et al., 2017) using bwa mem2 v2.2.1 (Vasimuddin, Misra, & Aluru, 2019). Resulting mapping files were converted to binary and sorted with SAMtools v1.19.2 (Danecek et al., 2021).

Genotyping for each accession was done with BCFtools v1.19 (Danecek et al., 2021), retaining only variant sites. Initial SNPs were filtered to remove low-quality genotypes, indels, and variants with high missing data or low minor allele frequency using VCFtools v0.16.1 (Danecek et al., 2021) with the options --max-missing 0.7 --min-alleles 2 --max-alleles 2 --minDP 5 --maxDP 200 --maf 0.05 --mac 3. Individuals with >50% missing data were excluded prior to pruning for linkage disequilibrium (LD) using PLINK v1.9 beta (Chang et al., 2015; Purcell and Chang, 2025) with a sliding window of 20 kb with a step size of 10 kb and the *r^2^*value of 0.5 calculated by PopLDdecay (Zhang et al., 2019).

### 2.5 Evaluation of genetic diversity

To identify potential duplicate accessions and closely related individuals, pairwise kinship coefficients were calculated using the KING-robust estimator (Manichaikul et al., 2010) implemented in PLINK v2 (Chang et al., 2015) on the LD-pruned dataset. Since KING kinship coefficients are scaled so that duplicate samples have kinship values close to 0.5, a less conservative range of ≈ 0.354-0.5 (the geometric mean of 0.5 and 0.25) was adopted to identify duplicates. Pairwise genetic distances were calculated in MEGA11 (Tamura, Stecher & Kumar, 2021) to provide a complementary measure of divergence between samples and support the identification of duplicate or highly similar accessions.

Genome-wide heterozygosity was estimated with ANGSD v0.940-dirty (Korneliussen, Albrechtsen & Nielsen, 2014) using the sorted BAM files, calculating the site allele frequency likelihoods (-dosaf 1) with the SAMtools genotype likelihood model (-gl 1). Only high quality sites were considered (-minMapQ 30 -minQ 20). The resulting site allele frequency likelihoods were used to estimate the maximum-likelihood site frequency spectrum with realSFS, and genome-wide heterozygosity was calculated as the proportion of heterozygous sites averaged over the genome.

We evaluated the inbreeding coefficient (*F*) with VCFtools v0.16.1 (Danecek et al., 2021) to explore how genetic diversity is maintained within and across farms and regions of Costa Rica. Given that cacao is typically an outcrossing species (Bartley, 2005) and farmers reported on their own breeding and seed-saving practices, inbreeding coefficients allow further investigation into the impacts of demographic history, gene flow, hybridization events, and human management on cacao diversity. Statistical comparisons of individual breeding coefficients among regions and farms were performed using Kruskal-Wallis tests (Kruskal and Wallis, 1952), followed by pairwise Wilcoxon rank-sum tests (Wilcoxon, 1945) with Benjamini-Hochberg correction for multiple testing (Benjamini and Hochberg, 1995). Linear models were fitted to evaluate regional and farm-level effects. All tests were conducted at a significance level of 0.05.

### 2.6 Population Structure

Population structure was assessed for the references plus Costa Rican samples using the final filtered VCF of 259 individuals, thinned to 111,884 SNPs (--thin 2500), and converted to Structure format using the populations module within Stacks v2.66 (Catchen et al., 2013). The Costa Rican only VCF file of 93 samples was thinned to 104,003 SNPs (--thin 2500).

Ancestral groupings were calculated using the sNMF algorithm implemented in the R package LEA v3.22.0 (Frichot & François, 2015; Frichot, François, & Gain, 2025). A range of ancestral clusters (*K*) from 1 to 25 were tested for both data sets, with a sparsity regularization parameter α = 100 and 500 iterations. The optimal number of *K* clusters was determined using cross-entropy values. Individuals with a maximum ancestry coefficient (MaxQ) of ≥ 0.8 were classified as pure lines. This cutoff was chosen to balance stringency and inclusivity and is consistent with thresholds previously applied in cacao population genomic studies (Argout et al., 2023; 85%; Todd et al., 2025; 70%).

Principal component analysis (PCA) was conducted using SNPRelate v1.44 (Zheng et al., 2012) and the LD pruned SNPs. The first four principal components were plotted to visualize patterns of clustering across all samples.

### 2.7 Phylogenetic analyses

To reconstruct evolutionary relationships among our Costa Rican collections, a maximum likelihood phylogeny was constructed using RAxML-NG v1.2.1 (Kozlov et al., 2019). The thinned VCF file containing 57,073 LD-pruned SNPs (--thin 5,000) was converted to FASTA using vcf2phylip (Ortiz, 2019). The GT+GTR genotypic model of evolution was used. Site concordance factors (sCF) were computed using IQ-TREE v2.2.2.7 (Minh et al., 2020) to provide a measure of support for each node.

We explored a phylogenetic network to visualize genetic relationships among admixed individuals and to identify clusters of closely related genotypes without assuming a bifurcating evolutionary history. Individuals exhibiting the greatest degree of admixture were identified using two complementary ancestry-based methods. First, Shannon entropy of ancestry coefficients were calculated to quantify the evenness of ancestry assignment across *K* clusters, with values normalized by the logarithm of the number of clusters, (log(*K*)) to produce a standardized metric between 0 and 1. Second, individuals were characterized by their maximum ancestry coefficient (MaxQ), representing the largest proportion of their genome assigned to a single cluster.

Consistent with our definition of pure lines (MaxQ ≥ 0.8), individuals with substantially lower dominant ancestry MaxQ (≤ 0.4) and high normalized entropy (≥ 0.65) were classified as highly admixed. The highly admixed subset, together with two representatives from each of the identified genetic clusters present within Costa Rica (total of 27 individuals) were was used to construct a phylogenetic network in SplitsTree v6.7.1 (Huson & Bryant, 2024) using the NeighborNet algorithm.

### 2.8 Landscape-level patterns of genetic variation

To examine how genetic variation in Costa Rican cacao is distributed and patterned across the landscape, we assessed isolation by distance (IBD) to quantify the effects of spatially-restricted gene flow among our collections (Van Strien, Holderegger & Van Heck, 2015). We quantified IBD by correlating Edward’s pairwise genetic distances between our Costa Rican collections with their corresponding geographic-based Euclidean distances. IBD was calculated among all collections and within each region individually (excluding Puntarenas with only two samples). Isolation by distance was inferred on individuals instead of populations since population groupings are relatively arbitrary and subjective based on habit, geographical distance, political boundaries, morphological differences, or cultural attributes (Manel et al., 2007; Pritchard et al., 2000). Specifically, we avoided the arbitrary grouping of a farmer’s population given the predicted exchange of genetic material between farms. Using the R package adegenet v2.1.11 (Jombart, 2008), the relationship between genetic and geographic distances was tested using a Mantel test with 10,000 permutations.

A hierarchical Analysis of Molecular Variance (AMOVA) was used to investigate how cacao populations in Costa Rica are defined and structured across the country, independent of inferred genetic groups. Farms were treated as the primary population unit, so that individual trees were nested within farms, and farms were nested within regions. The filtered SNPs were converted to a genind object and the AMOVA was performed using the poppr package v2.9.8 (Kamvar, Tabima, & Grünwald, 2014). Variance was partitioned among regions, farms within regions, and within farms, with variance components and THETA-statistics estimated using ade4 v1.7.23 (Dray and Dufour, 2007). Statistical significance was assessed using 999 permutations via Monte Carlo randomization. The AMOVA was repeated after excluding all samples from the Puntarenas region due to limited sampling.

To investigate genetic structure independently of sampling location, we conducted unsupervised clustering analyses at the individual level using a multivariate approach implemented in adegenet v2.1.11 (Jombart, 2008). Genotypes were first summarized using PCA to reduce dimensionality and account for linkage among loci. Genetic clusters were then inferred using a k-means clustering algorithm applied to retain principal components using the function find.clusters. The number of principal components (n.pca = 5) was chosen to retain the majority of genetic variation while minimizing overfitting. Clustering was then explored for K = 1 to 10, with K = 3 inferred as the levels of genetic differentiation to be visualized using discriminant analysis of principal components (DAPC). This approach maximizes among-cluster variation while minimizing within-cluster variation, allowing genetic structure to be evaluated independently of sampling information.

Lastly, we conducted a spatial AMOVA (SAMOVA) with geographic coordinates to identify geographically homogenous groups that maximize genetic differentiation among groups using SAMOVA v2.0 (Dupanloup, Schneider & Excoffier, 2002). SAMOVA assigns populations to a predefined number of groups (K) using a simulated annealing algorithm and molecular genetic distance plus geographic coordinates to maximize F_ct_, or the proportion of genetic variance explained among groups. Sampling locations (*i.e.*, farm) were retained as the basic unit rather than individual trees. Genotypes were converted to Arlequin format and the SAMOVA was run for K = 2 to 9, the greatest number of independent genetic groups identified by structure analyses. For each value of K, 10 independent simulated annealing runs were performed, each consisting of 20,000 iterations, to reduce the risk of convergence on local optima. The annealing process followed an exponential cooling schedule, where temperature was decreased multiplicatively at each iteration using a constant cooling factor (A = 0.9158), described by Dupanloup, Schneider & Excoffier (2002). Statistical support for inferred groupings was assessed using 1,000 permutations. The optimal number of groups was determined by comparing F_ct_ values across K and selecting the configuration that maximized among-group differentiation while maintaining geographic coherence.

## 3. Results

Sequences were obtained from 94 trees collected across 17 farms spanning four designated regions (Figure 1). Only one sample was excluded from downstream analyses due to negligible sequencing yield (<1Gbp). For the remaining samples, 4.7 to 62.7 Gbp of sequencing data were generated, with an average of 12.9 Gbp (Supplemental data S3). A total of 2,864,715 high-quality, unlinked SNPs were retained with an average depth of 34.06x (10.25-125.98x) and 4.31% missingness (0.14-48.39%; Supplemental data S4).

### 3.1 Evaluation of Genetic Diversity

KING-robust kinship estimation (Figure 2) revealed three pairs of duplicates (kinship coefficients > 0.354): GCM10-3 and GCM12-1; UPA1-3 and TAL9-1; and UPA6-2 and UPA6-3 - which were also duplicates of CCN-10 (kinship = 0.43l; Figure 2, red triangle). GTS5-8 was a clone of TSH-565 (0.46). Many individuals were also identified with a kinship range of 0.25-0.354, indicating first-degree kinship. GTS5-8 was closely related to references TSH-516 (0.35) and TSH-774 (0.29). Nine samples from Talamanca showed close kinship to the widely cultivated Matina1-6 clone (median kinship = 0.29; Supplemental data S5). Additional close relationships included TARS-31 with GTS2-3 (0.31) and various UF accessions (UF-613 with GTS2-4: 0.28; UF-168 and UF-676 with GCM11-1: 0.28).

**Figure 2.**
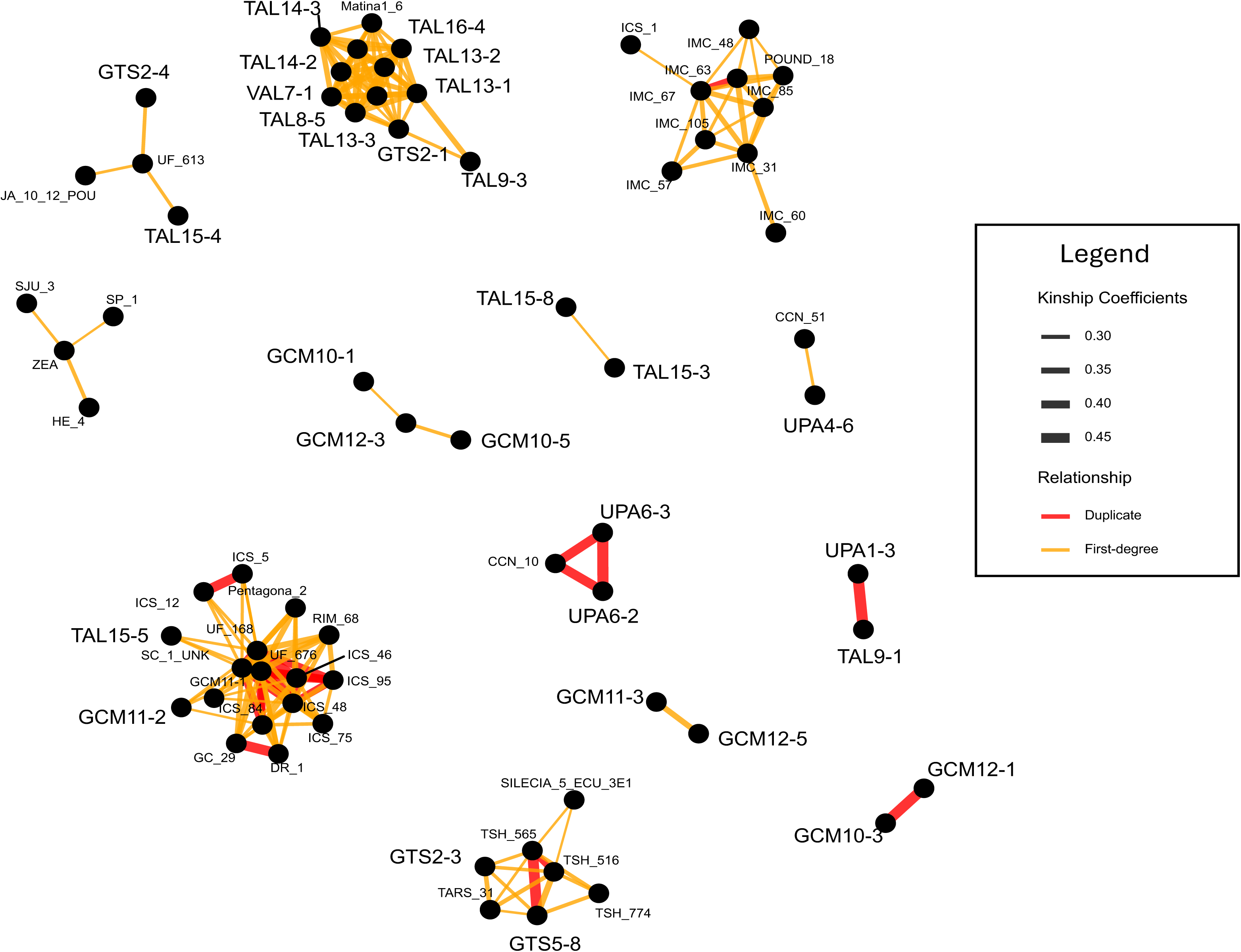
Kinship network graph depicting the genetic relationships of duplicate and closely-related Costa Rican individuals. Red lines connecting two collections indicate duplicate accessions, identified by kinship coefficients between 0.354 and 0.5. The thicker the line, the stronger the relationship. Orange lines indicate closely-related individuals, with first-degree relatives (parent-offspring, full siblings) determined by a threshold between 0.25 and 0.354, again with the thickness of the line indicating the strength of the relationship. The names of Costa Rican collections have been enlarged for emphasis. Clusters of closely-related reference accessions have been retained to demonstrate the general genetic similarity of investigated cacao genetic material.

Pairwise genetic distances confirmed most genetic relationships. Individuals exhibiting near-zero pairwise differences (< 0.005) were considered genetically indistinguishable. UPA6-2 and UPA6-3 were duplicates of CCN-10; GTS5-8 and TSH-565 were duplicates. All samples identified as Matina1-6 appeared as duplicates of CC-41. GCM11-3 and GCM12-5 were duplicates of Catongo, another common Amelonado variety. All genotyped Criollo reference accessions exhibited near-zero pairwise genetic differences, as did the three Costa Rican samples, GCM11-4, PUNT17-1, and the latter’s progeny, PUNT17-1F2 (Supplemental data S6).

Observed heterozygosity (H_o_) varied across individuals, farms and regions; ranging from 0.00068 to 0.01350 with an average of 0.00821 (Figure 3). Samples with extremely low heterozygosity, including one sample from Guácimo (H_o_ = 0.00068) and two from Puntarenas (H_o_ = 0.00069 and 0.00067), fell within the lower range observed for Criollo references (H_o_ = 0.000877 to 0.00466; Supplemental data S7). Inbreeding coefficients (*F*) for these samples were positive (*F* = 0.78 to 0.82), indicating excess homozygosity relative to Hardy-Weinberg expectations (Supplemental data S8). Inbreeding coefficients ranged from -0.57 to 0.82, with a negative mean (-0.11) and median (-0.21), indicating lower homozygosity than expected under random mating. Mean *F* differed significantly among farms (Kruskal-Wallis test, p < 0.001), with most farms having negative values consistent with predominantly outcrossing and substantial heterogeneity among samples. Observed heterozygosity also varied significantly across the sampled regions (Kruskal-Wallis rank sum test, p = 0.0039). Pairwise comparisons showed lower heterozygosity in Talamanca than in Guatuso-Upala (Wilcoxon rank sum test, p = 0.0041), while no significant differences were found between other regional pairs. In parallel, Talamanca trended towards less negative *F* values (Figure 3). Linear models attributed more variation in *F* to farm-level effects than region alone.

**Figure 3.**
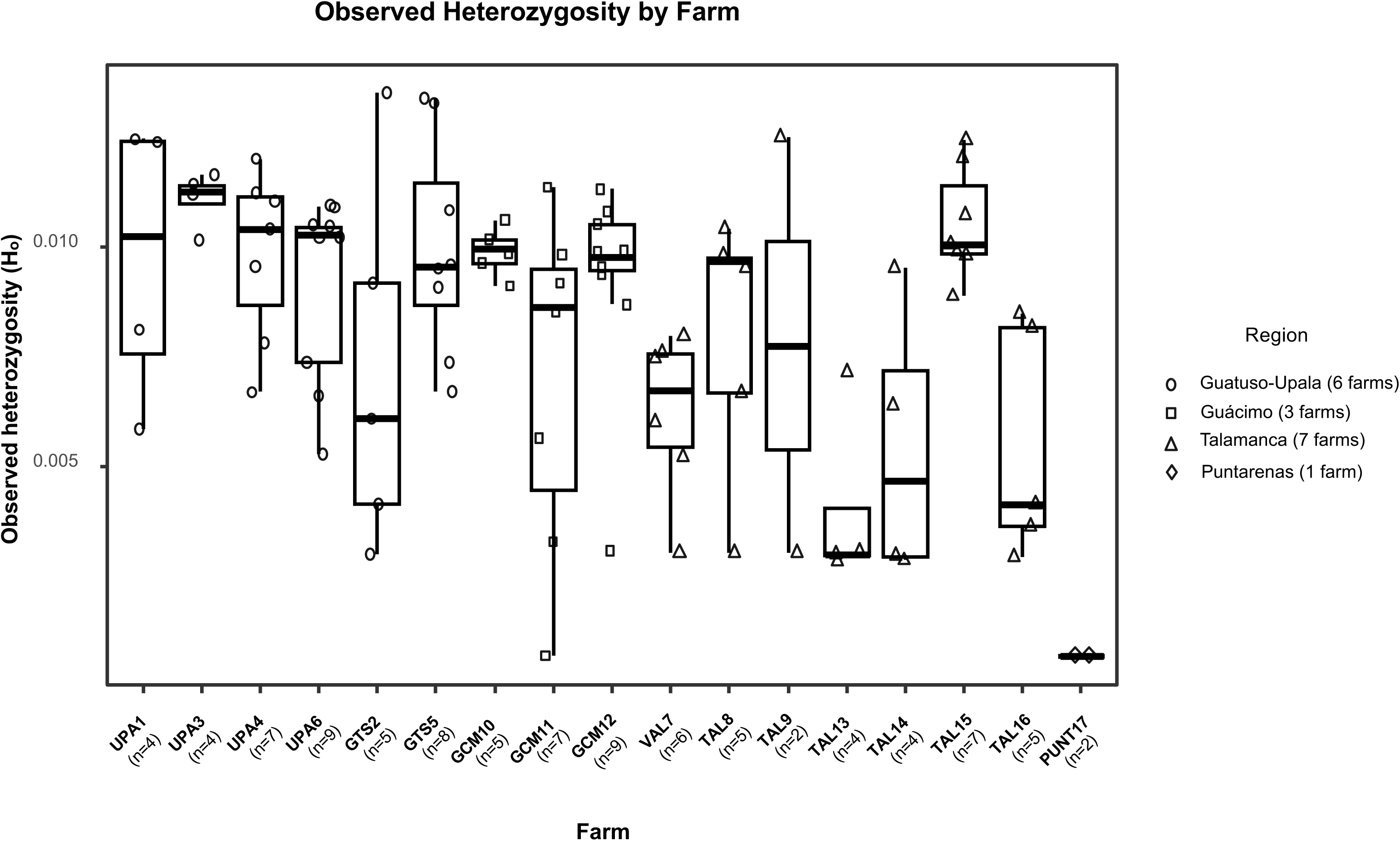
Observed heterozygosity was calculated using genotype likelihoods in ANGSD. Boxplots summarize the distribution of Hₒ within each farm (center line = median; box limits = interquartile range, whiskers = 1.5 x IQR). Individual trees are overlaid as points to show within-farm variation. Point shape denotes region of origin: circles = Guatuso-Upala, squares = Guácimo, triangles = Talamanca, and diamonds = Puntarenas. The number of trees sampled per farm is indicated below each farm as labelled on the x-axis.

### 3.2 Population structure

When all references were included, K = 17 provided the best fit for ancestral clusters, balancing resolution of established reference groups and patterns of diversity present within Costa Rica (Figure 4A). Outgroup *Theobroma* species clustered together (dark brown), as did Contamana (gray), Criollo (yellow) and Iquitos (teal). Three trees from Costa Rica were assigned to the Criollo group, GMC11-4 and PUNT171-F2 (pure lines) and PUNT17-1 (admixed; 75% composition). One individual, UPA4-6, was assigned to the Iquitos group (41% composition). Amelonado references (light green) were correctly identified, but appeared as admixed. For example, Matina1-6 had only 74% of its genome assigned to this ancestral cluster, with other previously characterized references ranging from 50-66% (median = 59%). In contrast, 14 Costa Rican accessions formed Amelonado pure lines (median = 90%, range = 81-98%; Supplemental data S9).

**Figure 4.**
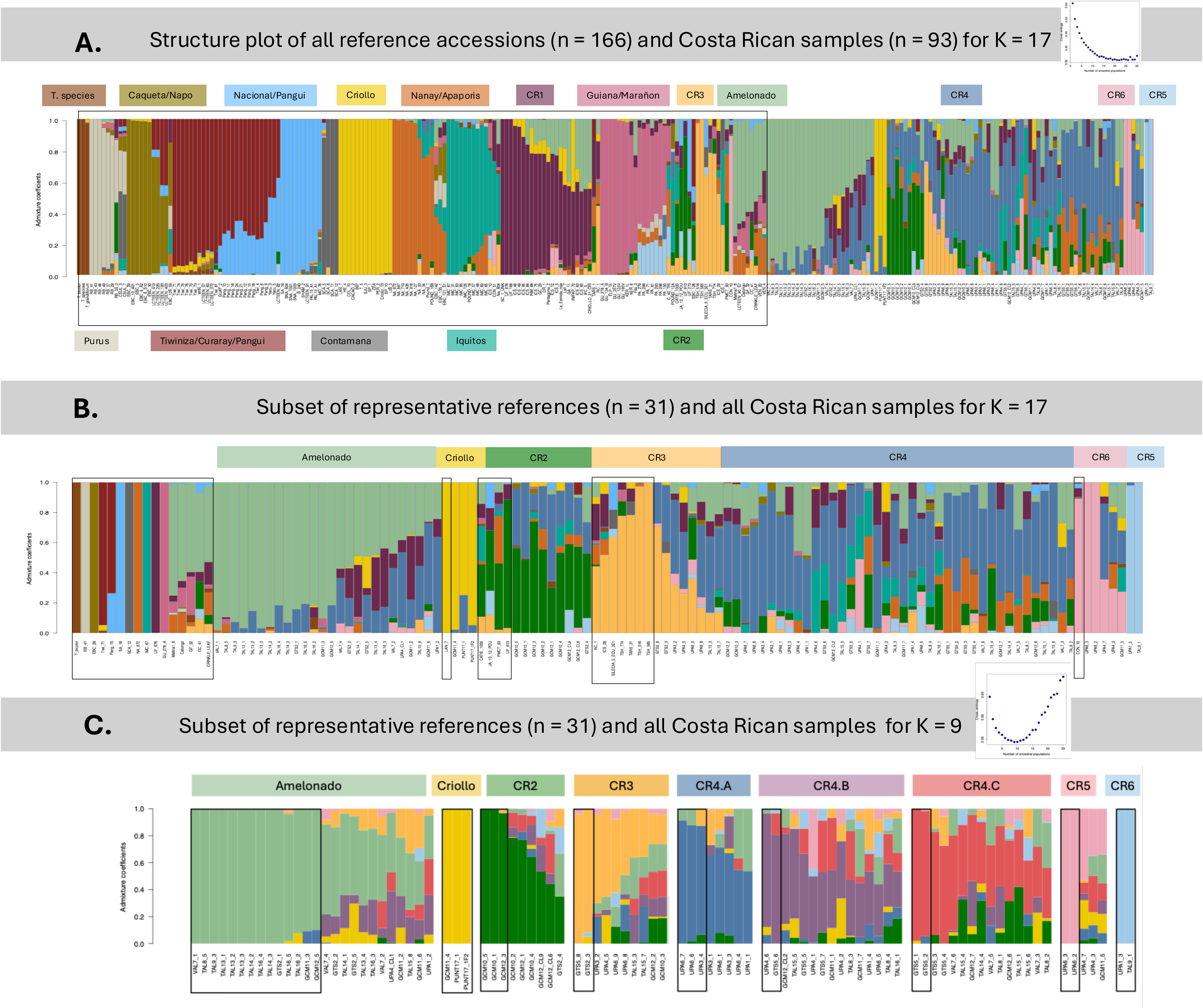
**(A)**. Structure plot showing inferred ancestry proportions at K = 19 for the full dataset, including all reference accessions and Costa Rican collections. Each vertical bar represents an individual, and colors respond to inferred genetic clusters. Reference accessions are grouped and labeled according to Motamayor et al. (2008), Fouet et al. (2022), and Argout et al. (2023) and are enclosed within a thin black box. This panel illustrates the placement of Costa Rican collections within the global genetic diversity of cacao. Previously uncharacterized genetic groups are labelled **CR1-CR6** (in order: dark purple, dark green, light orange, dark blue, light blue, light pink). **(B).** Structure plot as K = 19 showing a subset of panel A, including one representative reference accessions for each genetic group (left, enclosed in a thin black box) alongside all Costa Rican collections. Reference accessions clustering with Costa Rican collections have been repositioned to precede related materials and are also enclosed in thin black boxes. This arrangement highlights close relationships between our collections and known references groups, including Amelonado (light green) and Criollo (yellow) and specific reference accessions: CATIE-1000, JA-10-1-POU, PMCT-93, and UF-613 (**CR2**); RC-1, ICS-29, SILECIA-5- ECU-3E1, TSH-774, TARS-31, TSH-516, and TSH-565 (**CR3**); CCN-10 (**CR6**) **(C).** Structure plot for our Costa Rican cacao collections analyzed separately, inferred ancestral groups at K = 9. This analysis resolves the fine-scale population structure within the sampling, revealing pure ancestral groups previously detectable only as admixed components in panels A and B. Notably, **CR1** is no longer visible in this analysis. Individuals that previously clustered with reference accessions to form **CR2** (dark green) remain aligned with this group, with two individuals (GCM10-5 and GCM10-1) now resolving as pure lines. Similarly, trees that formed **CR3** (light orange) with reference accessions remain stable in this cluster, with GTS5-8 and GTS2-3 emerging as pure lines. CR4, which did not align with any known reference groups and appeared only as admixture in panels A and B, now solidifies into three distinct clusters, each containing pure individuals. Trees from Upala remain in **CR4-A** (dark blue). Two samples, UPA4-6 and GTS5-6 emerge as pure lines for a new group (**CR4-B**, soft purple) as do GTS5-1 and GTS2-2 for a third cluster (**CR4-C**, soft red). Previously identified groups, including Amelonado and Criollo, remain stable, across analyses, with14 individuals resolving as pure Amelonado and all three Criollo trees forming pure lines. Newly characterized groups from Costa Rican pure lines (**CR5** and **CR6)** also remain consistent.

While some reference ancestral groups were not recovered in our analyses, other groups that have not been characterized previously emerged as pure lines as early as K = 6 for United Fruit (UF) and Imperial College Selection (ICS) (Supplemental Figure 1). UF-676 and UF-168 appeared as pure lines of a distinct ancestral group (dark purple), along with SC-1-UNK. ICS accessions 46, 48, and 95 fell predominantly within this group (designated CR1) but included admixture with Criollo. Other ICS accessions were similarly assigned, although they showed significant admixture with Amelonado and Criollo. While only one Costa Rican individual (GCM11-2) was assigned to this group (33% composition), nearly 40% of the trees from our Costa Rican collections exhibited admixture from this group.

Other groups emerged that aligned with only one or a few reference accessions. UF-613 represented the only pure line of a previously uncharacterized ancestral lineage (dark green).

Accessions PMCT-93, JA-10-12-POU and CATIE-1000 were also assigned to this group, designated as CR2, but as admixed individuals. Trees collected from Guacimo (GCM) from farms 10 and 12 exhibit mixed ancestry with another uncharacterized lineage (CR4; dark blue). These individuals (GCM12-3, GCM10-5, GCM10-2, GCM12-CL6, GCM10-4) had 30-74% of their genome assigned to this ancestral group (median = 54%).

Trinidad Select Hybrid (TSH) accessions TSH-565 and TSH-615 formed their own ancestral group (CR3; light orange) despite their presumed hybrid origin (Figure 4A,B). Three admixed Costa Rican samples were assigned to this group, GTS5-8 (71%), GTS2-3 (53%) and UPA6-9 (36%). Although no references or Costa Rican collections were identified as pure lines of CR4, 46 of the sampled farmer materials were assigned to this group (median = 53%, range = 32-78%). No clear geographic patterns could be identified, as trees from every region except Puntarenas were represented within this ancestral group. However, this group was pervasive in admixture, present in 90% of our Costa Rican cacao collections (84/93).

A second group emerged with CCN-10 as the only pure line (CR5; light pink), clustering distinctly with two other pure accessions from Upala (UPA6-2 and UPA6-3; Figure 4A,B). Two other admixed individuals are assigned to this cluster, UPA4-1 and GCM11-5, with 47% and 27%, respectively.

One final group emerged in our analyses that did not align with any known references. UPA1-3 and TAL9-1 clustered together as pure lines (CR6; Figure 4A,B light blue). Evidence of admixture with this group was present in previously described Marañon (PA) accessions. CR6 was also present in individuals assigned to other ancestral groups. Notably, this ancestral group was present as admixture in ≈ 24% (22/93) of the Costa Rican collections and 16% (27/166) of the references.

When only Costa Rican collections were analyzed, K = 9 was the preferred number of ancestral populations (Figure 4C). Amelonado (14 individuals) and Criollo (three individuals) remained assigned to these groups. Both Criollo and Amelonado clusters were consistently distinguishable in the structure analyses from K = 3 to K = 9 (Supplemental Figure 2).

Most pure Amelonado individuals originated from the Talamanca region of Costa Rica, with three additional pure lines sampled from Guatuso and Guácimo. Three of these “pure” individuals had admixture from other groups, with TAL16-5 and TAL16-2 showing limited Criollo ancestry and GCM11-3 and GCM12-5 exhibiting ancestry from CR4-A (Figure 4C). The remaining Amelonado accessions included admixture from all other groups identified in the analysis, although GCM11-5 was the only individual with admixture from CR6 (Figure 4C).

The three pure Criollo individuals were collected from two farms, with two samples from a single farm in Puntarenas (PUNT17) and one from Guácimo (GCM11) (Figure 4C). Although several additional samples displayed notable Criollo ancestry (*e.g*., UPA4-2, 30%; GTS2-5, 30%; VAL7-3, 25%), ancestral contribution throughout the remaining samples was generally limited. While nearly 36% of Costa Rican collections (33/93) exhibited Criollo admixture, the majority of samples had minimal contribution (≤5% composition; Supplemental data S10).

Many of the novel ancestral patterns that emerged when comparing Costa Rican collections to reference accessions held, with the exception of CR1, which was largely composed of UF and ICS accessions and only two Costa Rican trees at K = 17. GCM11-2 and UPA4-2 both clustered with different groups at K=9, aligning with Amelonado and CR4-B, respectively (Figure 4C). CR2 remained stable. However, at K = 9, three accessions emerged as pure lines of this ancestral group, GCM10-5, GCM10-1, and GCM12-3. The remaining six individuals were all from Guacimo, except for one from Guatuso that was the most admixed of the group.

CR3 also remained largely the same. While no samples represented pure lines of this group at K = 17, two now separated as pure lines (GTS5-8 and GTS2-3). GCM12-3 and GCM10-3, which were identified as CR4 admixed individuals when references were included, joined CR3. CR4, which was previously visible as only admixed (dark blue in K = 17, Figure 4A,B), split into three distinct lineages, each with pure lines. Individuals with the highest CR4 ancestry (UPA6-6 and UPA6-7) remained, but appeared as pure lines (CR4-A; Figure 4C, dark blue). While many individuals with visible CR1 ancestry (*e.g.*, UPA4-6, TAL15-5, UPA4-2) resolved into the second newly emerged cluster, CR4-B (soft purple), others lacking the same background now cluster (*e.g*., UPA4-1, GTS5-7); suggesting CR4-B is not the same ancestry group as CR1, but a novel group (Figure 4C). Other samples that were assigned to CR4 with considerable admixture from Nanay lineages resolved into another Costa Rican cluster (CR4-C; Figure 4C, soft red). Many of the individuals within this group showed admixture with CR2 (Figure 4C) and Amelonado (Figure 4C).

Both CR5 and CR6 remained stable at K=9, with the same individuals. CR5 consisted of the two previously identified pure lines, UPA1-3 and TAL9-1, as did CR6 with its pure individuals UPA6-2 and UP6-3. The admixed individuals assigned to CR6 (UPA4-7, UPA4-3 and GCM11-5) demonstrated extensive admixture from all Costa Rican ancestry groups except CR5. Overall, persistence of unique, locally maintained germplasm derived from uncharacterized ancestral groups is clear. For the most part, Costa Rican farmer varieties are highly admixed.

Principal component analysis (PCA) revealed clustering patterns broadly consistent with ancestral groups. The first principal component (PC1) captured 13.4% of the total genetic variation, separating Criollo from all other samples, while PC2 (8.9%) further resolved non-Criollo clusters (Figure 5A, left). Amelonado individuals form a cohesive cluster, becoming less distinct in higher-order PCs (Figure 5A). In contrast, most Costa Rican collections are widely dispersed across genotype space. Samples that contain the same primary or majority genetic group tend to occur in close proximity within principle component space (Figure 5A), and admixed individuals exhibit considerable overlap.

**Figure 5.**
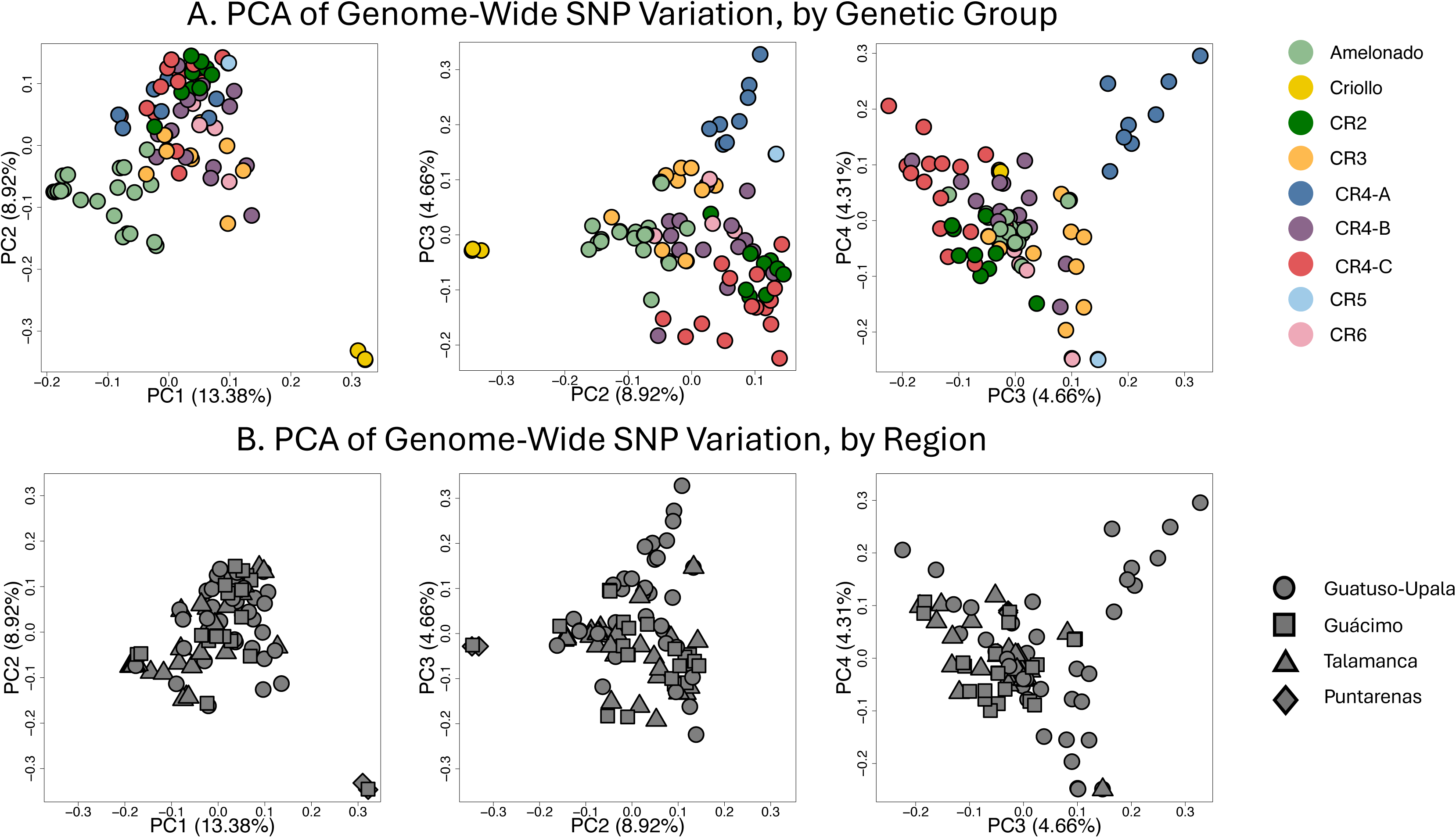
Principal component analysis (PCA) of Costa Rican cacao collections. **(A)** PCA plots showing PC1 vs. PC2, PC2 vs. PC3, and PC3 vs. PC4, with individual collections colored by inferred genetic groups. The variation explained by each PC can be found on each axis. **(B)** The same individuals plotted across the same principal component combinations, with shapes indicating region of origin. Guatuso-Upala is denoted by circles, Guácimo by squares, Talamanca by triangles, and Puntarenas by diamonds.

Patterns shifted when examining PC2 vs. PC3 (4.7%), with groups becoming more distinct (Figure 5A, middle panel): individuals of CR4-A separate from the core cluster of admixed samples, while CR4-C individuals cluster near the periphery. Criollo accessions remained distinct in PC2 vs. PC3. In PC3 vs PC4 (4.3%), Criollo individuals were no longer distinct, instead falling within the range of non-Criollo samples. Costa Rican genetic groups CR4-A, CR5, and CR6 begin to cluster. These patterns support finer-scale resolution captured primarily by higher-order principal components.

When samples were grouped by region (Figure 5B), clusters largely overlapped, with the exception of the two Criollo materials sampled in Puntarenas (Figure 5B, diamonds). No strong geographic patterns were observed across the remaining regions along PC1 vs PC2, however, subtle region-associated patterns emerged in higher order components, particularly within the Guatuso-Upala region (Figure 5B, middle and right panels). Together, these results suggest that broad-scale geographic structure is driven by deep lineage divergence, whereas local and region-associated structure among Costa Rican cacao becomes detectable only in higher-order axes of genetic variation.

### 3.3 Phylogenetic analyses

The maximum likelihood phylogeny (Figure 6) does provide some signal resolving individuals broadly into clades based on their dominant genotype (Figure 6). The three putative Criollo samples (Figure 6, node marked by a triangle) form a clade (sCF = 89.42%), with the two Puntarenas samples as a sister pair. Pure Amelonado individuals (less than 20% admixed) form a clade with 62 % sCF support (Figure 6, square symbol). All Costa Rican groups form clades except CR4-B, which is split across the phylogeny. UPA4-6 (sCF = 60.1%) is resolved in a more central position within the phylogeny, whereas GTS5-6 (sCF = 38.29%) occurs in a more distal clade (Figure 6, nodes marked by a circle).

**Figure 6.**
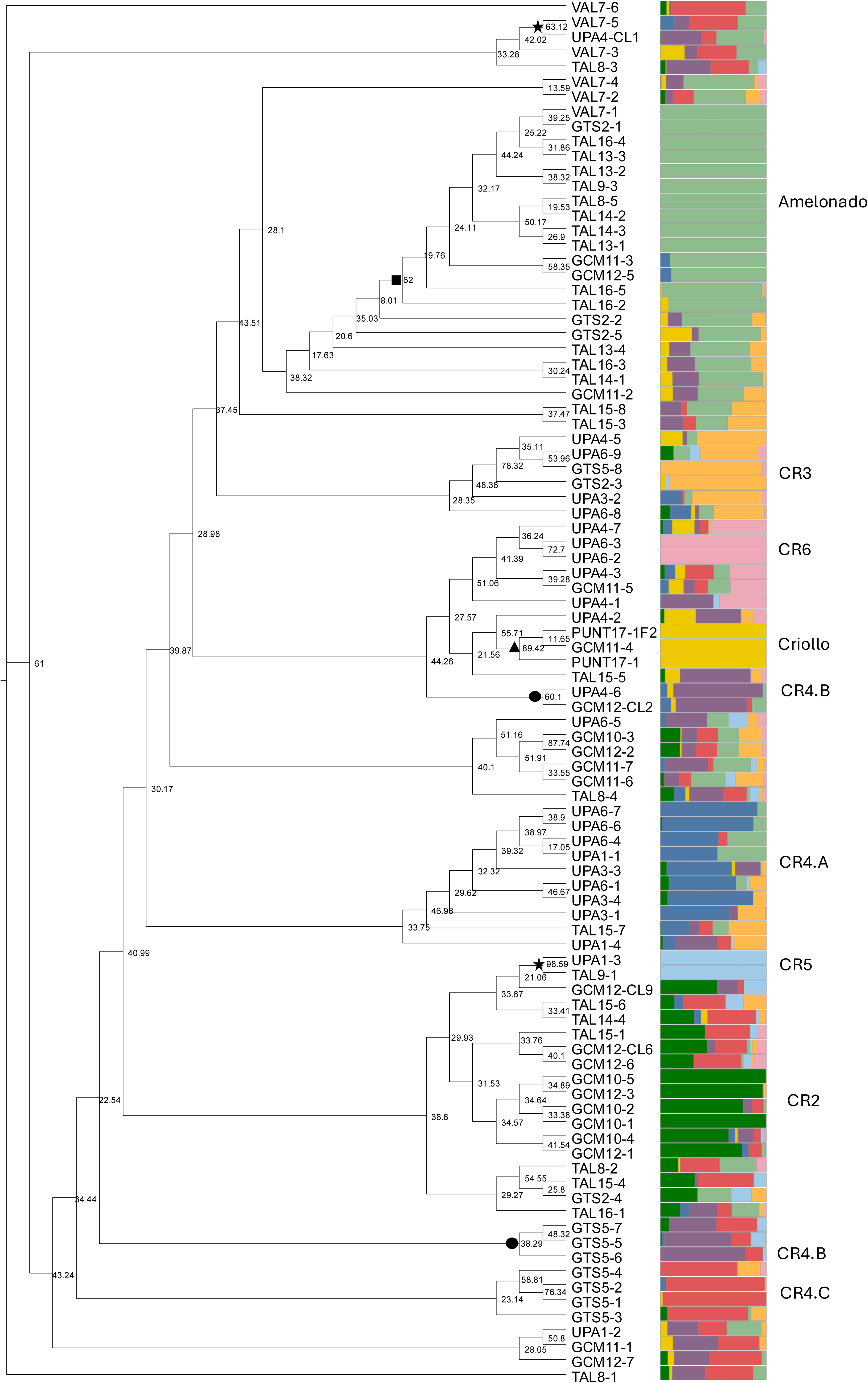
Maximum likelihood phylogeny of Costa Rican cacao from LD-pruned SNPs. Individual collections are shown at the tips, with the structure bar plot for K = 9 corresponding to the same individuals included in the phylogeny. Ancestral lineages are also labeled. Support values at the nodes represent site concordance factors. Starred nodes highlight sister pairs composed of individuals from geographically distant regions (Guatuso-Upala and Talamanca).

Closely related individuals were frequently collected from the same farm; however, we observed no consistent regional patterns. For instance, materials from Upala, the northernmost sampling on the border with Nicaragua, formed sister pairs with materials from Talamanca in the South Caribbean region of the country (*e.g.,* UPA4-CL1 and VAL7-5; UPA1-3 and TAL9-1, starred nodes)

The phylogenetic network incorporating 29 individuals (Figure 7) recapitulated several of the genetic relationships identified in the population structure and phylogenetic analyses. The Criollo collections (Figure 7; GCM11-4 and PUNT17-1, yellow) showed strong separation. Amelonado accessions also form a distinct cluster (Figure 7; VAL7-1 and TAL8-5, light green). Most novel groups represented by pure individuals formed their own groups or closely connected tips (Figure 7; CR2, dark green; CR4-A, dark blue; CR4-C, soft red; CR5, light blue; CR6, light pink). Several individuals were found to occupy intermediate positions between major genetic groups; *e.g.*, TAL8-4, which exhibits admixture from every group, was situated between CR5 and CR6. Likewise, UPA6-5 is positioned closest to the CR4-B pure lines, which is its dominant ancestral group based on structure analyses (Figure 4C, third individual from the right within the CR4-B group). The same is true for TAL16-1, which while belonging to the CR4-B group, had high Amelonado admixture and was resolved as clustering between CR4-B and Amelonado (Figure 4C, last individual within the CR4-B group).

**Figure 7.**
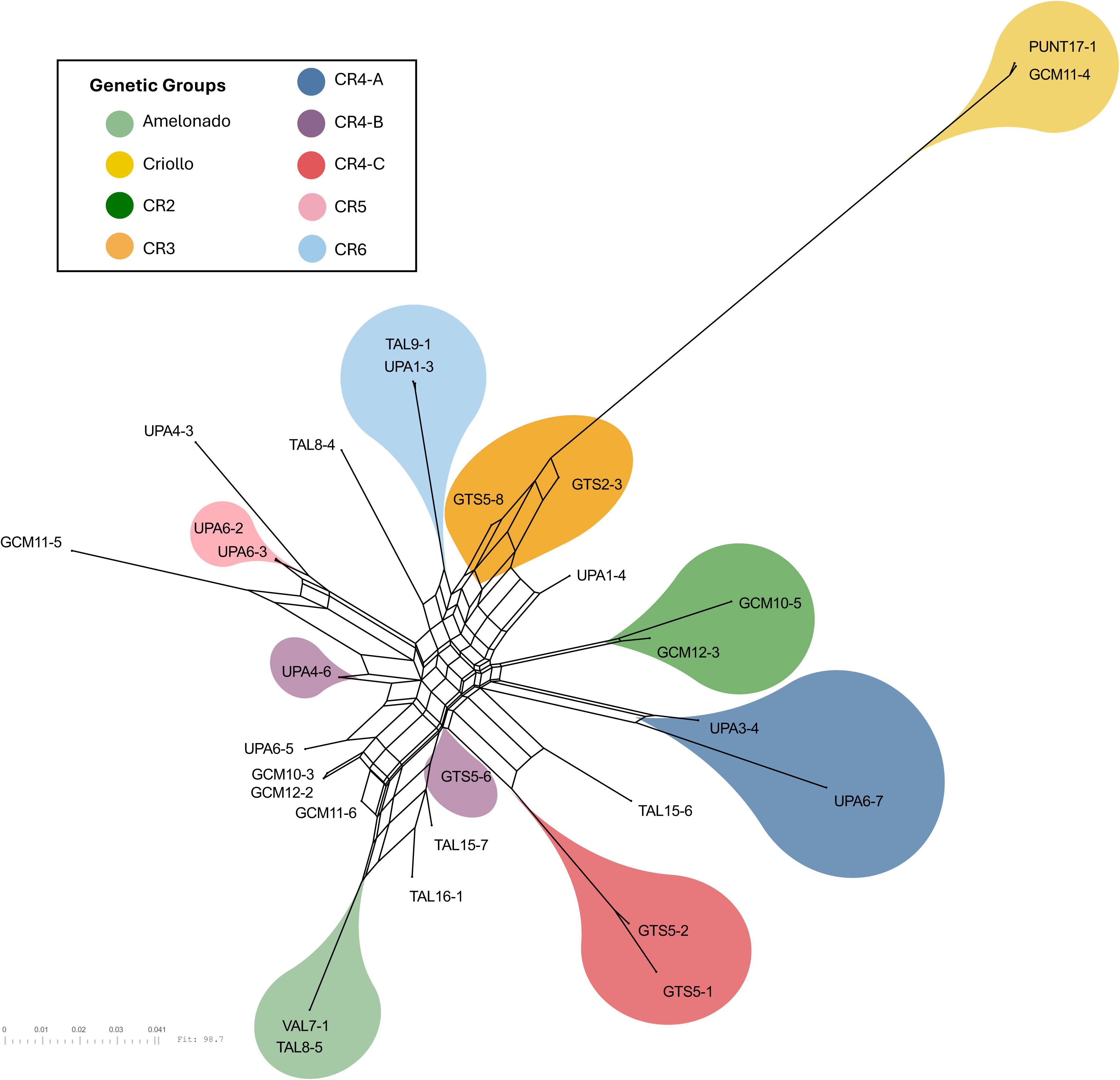
Phylogenetic network inferred using the NeighborNet algorithm. The network is composed of 11 of the most admixed samples (MaxQ ≤ 0.4 and normalized entropy ≥ 0.65) alongside two representatives per each of the nine genetic clusters present in Costa Rica (colored according to their group). In the NeighborNet network, each tip represents an individual sample. Edge lengths are proportional to split weights derived from the genetic distance matrix and reflect the strength of separation between samples. Parallelogram-like structures indicate incompatible or conflicting splits in the data that are not strictly bifurcating. Individuals positioned within reticulated regions (the parallelogram-like structures) or between well-defined clusters may reflect conflicting phylogenetic signals, which can arise from processes such as admixture, gene flow, incomplete lineage sorting, or recombination.

### 3.4 Landscape-level patterns of genetic variation

Isolation by distance revealed no significant correlation between pairwise genetic distance and geographic distance across the country (*r =* 0.05, *p* = 0.11; Figure 8A). Similar patterns were observed at the regional scale; separate Mantel tests within Guatuso-Upala, Guácimo, and Talamanca showed no significant correlation between geographic and genetic distance (all *p* > 0.25). Correlation coefficients remained weak (Guatuso-Upala: *r* = 0.04; Guácimo: *r* = -0.13; Talamanca: *r* = -0.05), further supporting the absence of spatially structured genetic variation.

**Figure 8.**
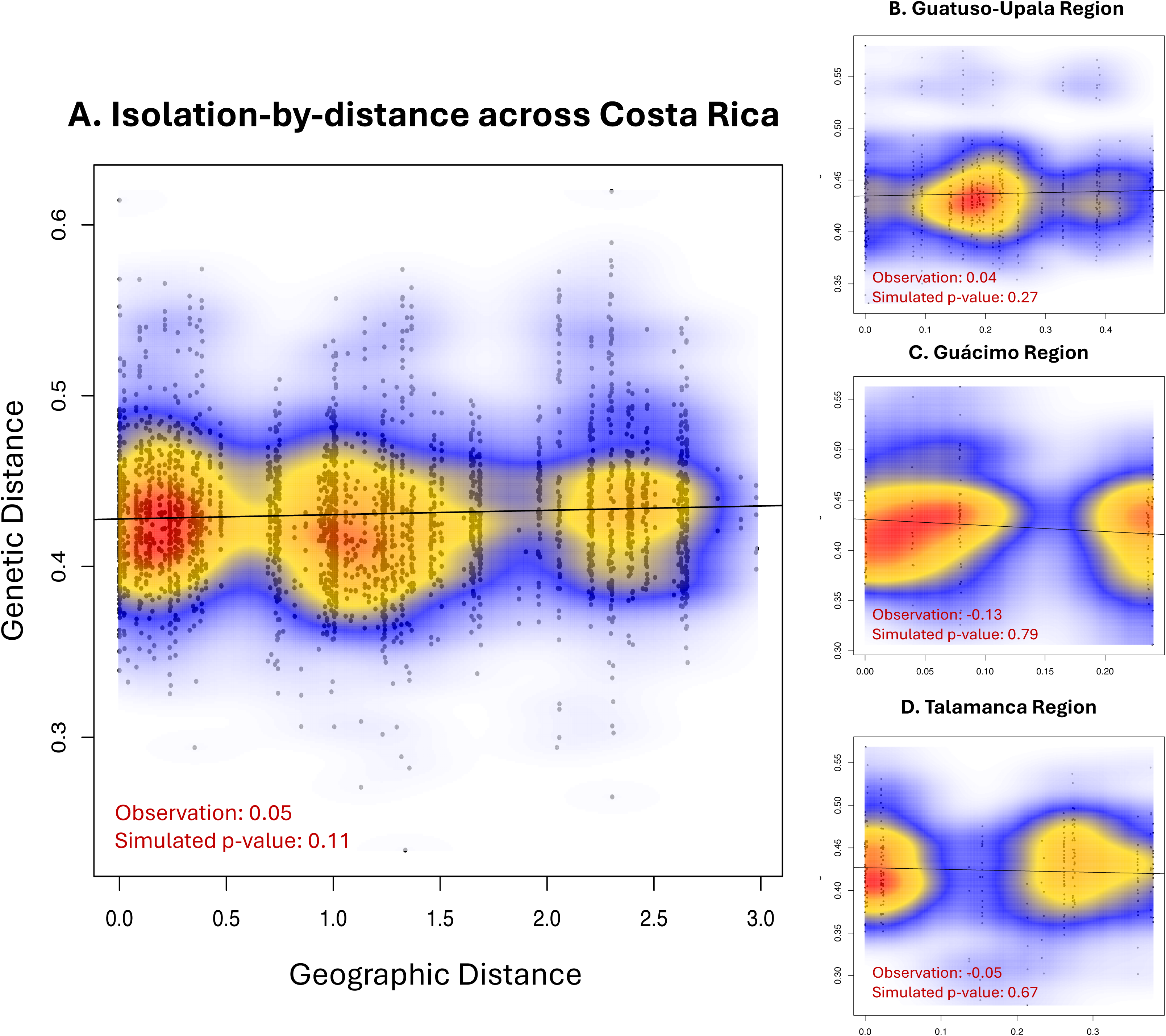
Results of Isolation by Distance (IBD) analyses among Costa Rican samples. Panel A shows the overall relationship between genetic and geographic distance across the whole country, with an observed Mantel correlation of 0.05 (p = 0.11), indicating no significant IBD at the country scale. Panels B, C, and D show IBD within the regions with extensive sampling, ordered from north to south. Observed correlations and resulting p values also demonstrate a lack of significant isolation by distance at the regional scale.

AMOVA confirmed that most genetic variation occurred within farms (82-88%), with smaller but significant components among farms within regions (6-7%) and among regions (5-11%). When samples from Puntarenas were excluded, the proportion of variation attributed to regional differentiation decreased by nearly half, indicating that this region contributes disproportionately to the observed spatial genetic structure. In contrast, differentiation among farms within regions remained stable across analyses. Global F-statistics corroborated these patterns, with a low but significant overall F_st_ (0.079) and near-zero F_ist_ (0.0009), reflecting weak population differentiation and limited inbreeding.

Pairwise F_st_ among farms were highly variable, (-0.12 to 0.79), emphasizing substantial heterogeneity across sampling locations. Notably, farms containing Criollo or Amelonado individuals showed the highest pairwise F_st_ values relative to other farms (PUNT17 with Criollo ≈ 0.55; TAL13 with Amelonado ≈ 0.20). While the AMOVA and pairwise F_st_ analyses revealed measurable genetic differentiation among farms and regions, on a locus-by-locus basis, this differentiation was not statistically significant (g.star for all loci = 0, *p* = 1). Although certain farms exhibited distinct allele frequencies at the population level, most loci did not show strong differentiation among farms.

As expected, Criollo genotypes formed a distinct cluster in the unsupervised clustering (DAPC) analysis (Figure 9). A second cluster was composed of Amelonado individuals and those with high Amelonado ancestry. The third cluster included all remaining samples,underscoring the difficulty of assigning individuals to discrete populations. These patterns reinforce the mosaic of genetic ancestries within Costa Rica, with clear differentiation only for Criollo and Amelonado individuals.

**Figure 9.**
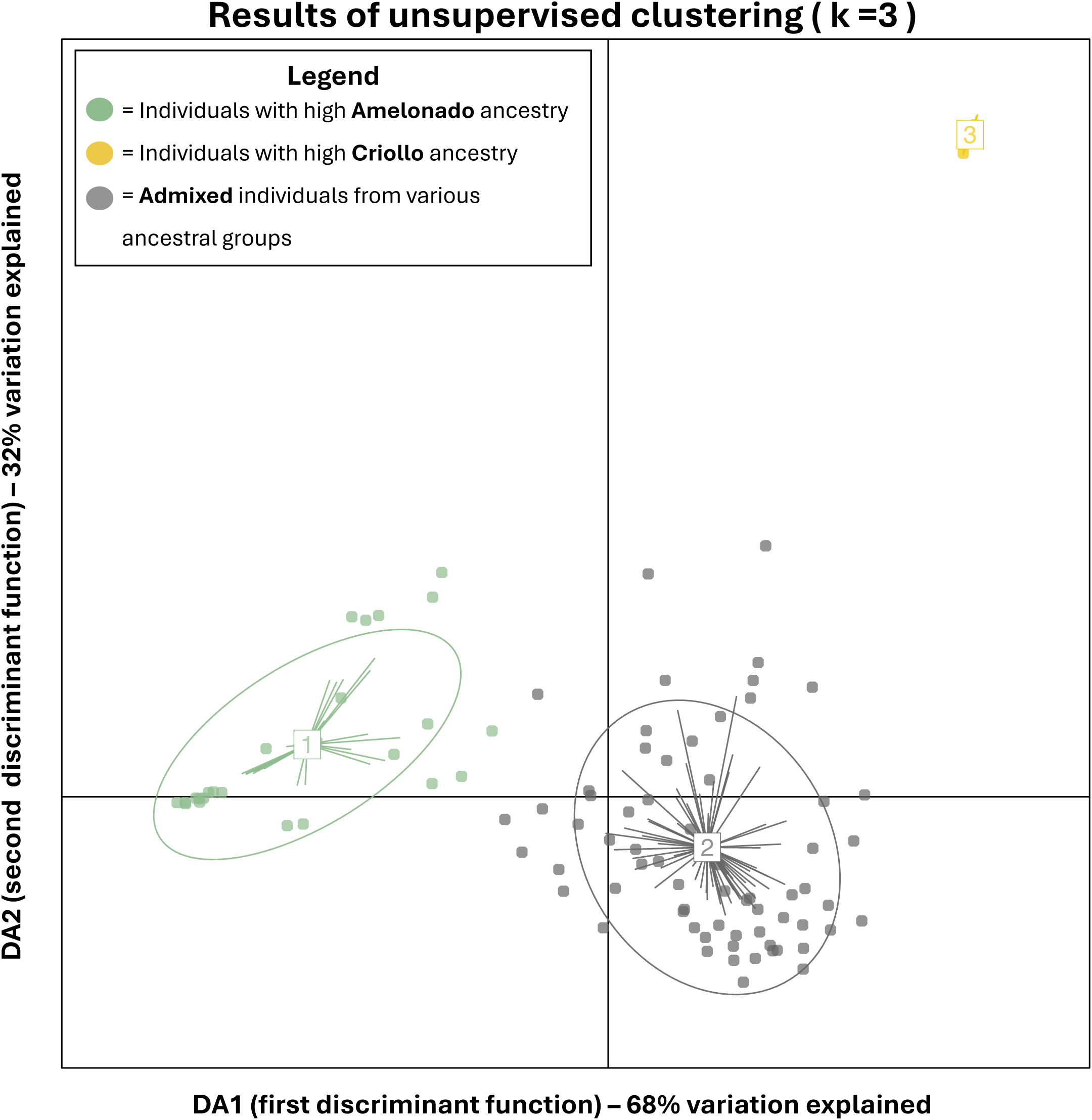
Discriminant Analysis of Principal Components (DAPC) of Costa Ricana cacao genotypes, showing genetic clustering of individuals independent of sampling location. Each point represents an individual tree, with colors corresponding to unsupervised genetic clusters. Cluster 1 (light green) corresponds to Amelonado materials and those with dominant Amelonado ancestry, while Cluster 3 corresponds to Criollo materials. Cluster 2 (dark grey) reflects individuals with mixed ancestry. Axes represent the first (DA1) and second (DA2) discriminant functions, explaining 68% and 32% of the between-cluster genetic variation, respectively. The spatial distribution of points and clusters highlights the mosaic of genetic ancestries across Costa Rican cacao collections, illustrating that while Criollo and Amelonado individuals are genetically distinct, most trees collected exhibit extensive admixture.

SAMOVA indicated that the optimal number of groups was K = 2, explaining 24.2% of the genetic variation. Comparatively, from K = 3 to 9, the average F_cts_ remained around 10% (Supplemental data S12). At K = 2, only one farm (GCM12) was assigned a unique group, with all other farms clustering together. Nine trees were sampled from this location, making it one of the most intensely sampled locations, matched only by one other farm in Upala. Individuals from GCM12 were largely admixed, with a single pure tree assigned to the Amelonado group, and another to the CR2 group. The remaining individuals were assigned to each of the other Costa Rican clusters identified, except for Criollo, CR5, and CR6 - highlighting the high genetic diversity at this farm. Although this pattern suggests some degree of population differentiation, the isolation of a single farm may reflect SAMOVA’s sensitivity to localized allele frequency differences rather than a strong, biologically meaningful separation.

## 4. Discussion

This study reveals the presence of novel ancestral groups in Costa Rican cacao, contributing to extensive admixture across the collected individuals and within the sampled farms. Of the 94 collections analyzed, three can be confidently assigned to Criollo, evident by their distinct clustering across all analyses. Although rare, their presence confirms that true Ancient Criollo germplasm persists within farmer-managed systems. In contrast to these few Criollo trees, the dominant pattern across Costa Rican cacao is extensive genetic admixture. Sixty trees collected and analyzed reflect mixed ancestry from known genetic groups represented in reference collections and from previously uncharacterized ancestral lineages. Several locally consistent groups (CR2, CR4-A, CR4-B, CR4-C, and CR5) were recovered, revealing regionally unique or historically under-sampled diversity. Related to widespread admixture, levels of observed heterozygosity vary drastically; ranging from nearly homozygous Criollo genotypes to highly heterozygous individuals consistent with ongoing outcrossing. The combination of high admixture and heterogeneous diversity highlight complex histories of introduction, recombination, and local propagation rather than uniform breeding or replacement by clonal materials. Similar patterns of genetic complexity have been observed in other studies of Mesoamerican cacao, where multi-wave introductions and farmer selections have produced highly admixed populations (Motilal et al., 2010; Ji et al., 2012)

Neither phylogenetic nor spatial analyses reveal strong geographic structuring across the collected Costa Rican cacao individuals. Phylogenetic relationships confirm extensive intermixing of trees and germplasm from different regions, while IBD hierarchical AMOVA, and SAMOVA detect no consistent associations between genetic and geographic distance. By identifying the existing diversity conserved within Costa Rica, we found that the current genetic landscape of cultivated cacao does not align with existing reference frameworks (Motamayor et al., 2008; Foet et al., 2022; Argout et al., 2023). Rather, we identified a complex tapestry of genotypes within diversified production systems, in which farmers play a central role in shaping and maintaining dynamic cacao diversity.

### 4.1 Value of Criollo germplasm

The identification of Ancient Criollo in Costa Rica is notable. The greater prevalence of Criollo-admixed individuals is consistent with historical introgressions and the widespread replacement of Criollo by more productive materials (Lachenaud and Motamayor, 2017). While we included some collections from Indigenous territories, which are frequently underrepresented in genomic studies and likely harbor unique diversity (Schulz, Becker & Götsch, 1994; Hernandez Marentes et al., 2022; Sterling et al., 2025), future efforts should prioritize more extensive sampling. Criollo cacao likely persists elsewhere in the region, often sought for its historic and cultural significance, superior quality, and association with fine flavor (Motilal et al., 2010; Maharaj et al., 2019; López et al., 2021). Costa Rica has long prioritized the production of local, quality cocoa over bulk production and was one of only 10 countries recognized by the International Cocoa Organization in 2016 as exporting 100% fine flavor cocoa (International Cocoa Organization, 2016). The detection of both pure Criollo and Criollo-admixed materials suggest that fine flavor-associated ancestry remains embedded within contemporary production systems.

### 4.2 Identifying unique Costa Rican cacao germplasm

Costa Rican cacao exhibits extensive admixture, with no clear geographic patterns of population differentiation; supporting our conclusion that gene flow is largely human-mediated, rather than governed by distance or landscape features. As a high-value crop, farmers frequently exchange planting material (Lagneaux et al., 2021), save seeds and clonally propagate elite genetic lines (Abdul-Karim et al., 2023), resulting in high genetic connectivity. The identification of the same genotype in both Upala and Talamanca - at opposite ends of the country- provides a clear example of human-mediated gene flow.

The identification of locally consistent yet uncharacterized genetic ancestry groups underscores the importance of geographically inclusive sampling. Studies with expanded sampling in other cacao-producing regions have revealed uncharacterized lineages (Zhang et al., 2012; Foet et al., 2022; Argout et al., 2023). Our results illustrate that the number and boundaries of cacao genetic groups become refined as geographically localized diversity is more thoroughly documented. The relative scarcity of genomic resources relevant to Costa Rica, and Central America more broadly, contributes to this complexity. Although the availability of whole genome sequencing data has increased since 2018 for cacao, most sequenced accessions originate from South America, particularly the Amazonian center of origin (Fouet et al., 2022; Argout et al., 2023). Characterizing the true genetic diversity of Costa Rica and Central America more broadly requires additional sampling to resolve the complexity of cacao.

### 4.3. The complexity of global cacao population structure

Global cacao genetic diversity is most often discussed in relation to the 10 major genetic groups identified by Motamayor et al. (2008), which serves as the primary framework for interpreting population structure. Although subsequent studies have proposed further subdivision (Thomas et al., 2012; Gutíerrez et al., 2021), these revisions remain contested (Todd et al., 2025), reflecting the fluid boundaries among groups and the inherent difficulty of resolving discrete lineages in a system shaped by admixture, gene flow, and uneven sampling. The absence of strong clustering within established genetic groups, regardless of sequencing methods, reference genome, reference panels, sequencing depth suggests that much of the observed diversity does not align with current references.

Several previously characterized ancestral groups emerged differently in our analyses than described previously. Lineages defined as distinct and pure ancestral populations (Argout et al., 2023) appeared as admixed or clustering with other ancestral groups: some Napo accessions (LCTEEN-413 and LCTEEN-220) appeared indistinguishable from Caqueta (Figure 4A; dark gold), others (LCTEEN-193 and LCTEEN-403) showed admixture from Twiniza (Figure 4A; dark red) and Nacional (bright blue); Apaporis (EBC) and Marañon (PA) accessions are heavily admixed with ancestry components spanning multiple groups (Figure 4A; CR6, light blue).

Reference accessions of United Fruit (UF), Imperial College Selection (ICS), Trinidad Select Hybrids (TSH), and Colección Castro Naranjal (CCN), described as Trinitario and hybrids (Todd et al., 2025), cluster as CR1, CR2, CR3, and CR5. These results highlight the difficulty of resolving discrete ancestral lineages when admixture is pervasive; suggesting that global cacao structure is more reticulate than strictly hierarchical, particularly when reference panels are broadened to include previously underrepresented diversity.

In perennial fruit species, long domestication histories, repeated hybridization, and introgression from multiple progenitors frequently obscure clear genetic boundaries (Zhuang et al., 2023). In apple, extensive gene flow from wild relatives produce weak and diffuse population structure (Cornille et al., 2012), while human-mediated movement and clonal propagation in grapes blur genetic clusters, challenging the assignment of discrete ancestral groups (Myels et al., 2011). In apricot, multiple geographically-isolated diversification and domestication events resulted in gene flow (Liu and Cornille et al., 2019). Biologically relevant ancestral groups tend to correspond to geographical regions, a pattern in cacao when sampling is sufficiently broad (Liu and Cornille et al., 2019; Fouet et al., 2022; Argout et al., 2023).

Complex admixture does not preclude biological interpretation, but rather necessitates careful evaluation of robust clusters that are geographically coherent. While the major genetic groups of cacao remain an essential reference framework, composition shifts as new regions are sampled and genomic resources are expanded. The emergence of novel Costa Rican genetic groups should not be interpreted as contradictions to previous studies, but as evidence that cacao population structure is dynamic and sensitive to sampling breadth, marker choice, and analytical approach (Schwartz and McKelvey, 2009; Shringarpure and Xing, 2014; Linck and Battey, 2019).

Given the lack of Costa Rican-relevant genomic resources and the unresolved ancestral lineages, future research needs to target additional sampling from local farmer varieties, especially in Puntarenas and the South Pacific of the country, where representation is severely lacking. Additional sampling from other Central American countries may illuminate patterns of genetic diversity at a broader scale. Cacao farms within these countries, and even within national genebanks, could be the origin of novel ancestral lineages. However, the characterization of these materials and collections are crucial if regional cacao diversity is to be understood. In a study of six Latin American genebanks, in all countries except Honduras, more than half of the collections had not been characterized or evaluated (Cecarelli et al., 2022). While many germplasm collections are understaffed and underfunded, the characterization of genetic resources ultimately underpins our ability to understand, utilize, and effectively conserve biodiversity (Fu, 2017; Engels, Ebert & van Hintum, 2024).

### 4.4 Implications for future resiliency

Additional collections and genomic characterization of cacao diversity within Central America and across all cacao-producing countries will be critical for meeting future production challenges. Under future climate predictions, harnessing currently underrepresented adaptive genetic variation in elite breeding materials and germplasm collections will become a priority.

Traditional landraces and farmer varieties harbor diversity shaped by long-term farmer-crop-environmental interactions that confer local adaptation, stress resistance, disease resistance, and nutritional value - making them invaluable for future crop improvement and sustainability (Dias, Anuruddi & Fonseka, 2024). Furthermore, while modern cultivars are superior in yield and uniformity, they have undergone intense selection resulting in narrower genetic diversity (Casañas et al., 2017). Our work suggests underlying cultural attachments and preferences among farmers for maintaining local crop diversity, increasing resiliency under future climate scenarios (Arias Montevechio et al., 2023; Lazaridi et al., 2024)

Emerging genomic approaches such as genomic-offset and genotype-environment association are powerful tools for predicting responses to future climates and identifying genotypes that are pre-adapted to future environmental extremes. These methods have been successfully applied in other crop systems to quantify the genomic change required for local adaptation, supporting local germplasm for minimizing maladaptation and creating resiliency under extreme climate scenarios (McLaughlin et al., 2025). In this context, the systematic characterization of diverse cacao materials, from farms and undercharacterized germplasm collections, may reveal alleles and polygenic combinations associated with tolerance to heat, drought, disease pressure, and other biotic and abiotic stresses that are likely to become more severe with climate change.

## 5. Conclusion

As interest in strengthening Costa Rica’s domestic chocolate industry continues to grow, a clear understanding of the genetic composition of farmer-managed genetic diversity is valuable. Our results suggest the farmer-maintained cacao harbors genetic diversity distinct from current reference accessions, potentially representing local populations and individuals with Criollo ancestry. These findings highlight opportunities for conservation and further characterization of germplasm not currently included in global collections, while deepening our understanding of the cacao diversity underlying the fine flavor chocolate produced in Costa Rica. As additional cacao accessions from Costa Rica and Central America, more broadly, are sequenced, we expect to refine local genetic clusters, identify additional Ancient or Modern Criollos, and further resolve patterns of admixture and regional genetic structure.

## Supporting information

Supplemental Data with all sheets

## Acknowledgements

The authors would like to thank Ruth M. Castro-Vásquez for providing laboratory space and resources in Costa Rica during sampling. Many thanks to Jen Grenier and the Cornell Genomics Innovation Hub, as well as Linda Cote and the Cornell Genomics Facility for assistance with library preparation and sequencing. The authors would also like to thank the Cornell BioHPC for computational resources. Many thanks to Pavel Dimens for his helpful suggestions on additional analyses, The research was partially funded by The Regional Fund of Agricultural Research and Technological Development (FONTAGRO) as part of the project ATN/RF 20629-RG.

## Data Accessibility

Newly generated sequencing data has been submitted to the Sequence Read Archive with accession numbers provided in Supplemental Table S1.

## Author contributions

CEH, EMH, MAG, JBL, HAT and CDS conceptualized the study; JEG, CEH and EMH developed sampling methodology; EMH and LS collected samples; EMH and JBL generated the sequencing data; EMH analyzed the data; EMH and JBL wrote the first version of the manuscript; EMH, JBL, MAG, HAT, and CDS revised the manuscript and all authors approved the final version.

